# Self-healing, biocompatible bioinks from self-assembled peptide and alginate hybrid hydrogels

**DOI:** 10.1101/2024.06.19.599807

**Authors:** Emily H. Field, Julian Ratcliffe, Chad J. Johnson, Katrina J. Binger, Nicholas P. Reynolds

## Abstract

There is a pressing need for new biomaterials that are printable, stiff and highly biocompatible. This is primarily due to the inverse relationship between the printability and viscosity of hydrogels. Cell-laden, printable, rigid biomaterials are needed for replicating stiffer tissues such as cartilage in regenerative medicine, modelling the fibrosis of tissue and cancer microenvironments, as well as in non-cellular research fields such as biosensors. Here, we have designed a hybrid material compromised of self-assembled Fmoc-FF peptide assemblies dispersed throughout a sodium alginate matrix. The resultant hybrid bioink has a stiffness up to 10 times greater than sodium alginate alone but remains highly printable, even when laden with high concentrations of cells. In addition, the thixotropic self-assembled peptide assemblies gave the hybrid bioinks highly desirable self-healing capabilities. The choice of solvent used to initially dissolve the peptides made significant differences to both the physical properties and the biocompatibility of the bioinks, with the best performing able to support the growth of encapsulated macrophages over 5 days. Our developed hybrid materials allow the bioprinting of materials previously considered too stiff to extrude without causing shear induced cytotoxicity with applications in tissue engineering and biosensing.

## 2. Introduction

3D bioprinting is rapidly becoming the gold standard for creating artificial tissue constructs due to its accuracy and ability to create complex 3D shapes and patterns [1]. Extrusion based bioprinting allows the printing of ‘living tissues’ where biomaterials laden with cells are printed into specific shapes in tailored bioinks [2-4]. For example, such cell laden materials have been used to create artificial 3D cartilage and muscle tissues with environment specific cells and constituents [5-7]. Extrusion-based bioprinting is also being used for non-cellular applications like precisely printing electrode interfaces on biosensors to create bio-electronic materials [8, 9].

Viscosity and printability tend to have an inverse relationship, making the printing of rigid materials challenging [10, 11]. Rigid materials are necessary for cartilage (Youngs modulus, 100-1000 kPa) and bone engineering (osteoid 35 kPa, initial calcified matrix 150 kPa), mimicking the fibrosis of tissue (4-25 kPa), long-term wound healing gels and prolonged drug delivery systems [12-18]. However, cell-laden materials must be printed at extrusion pressures that are sufficiently low so as not to damage encapsulated cells. Commonly used materials for tissue engineering and bioprinting include natural polymers such as hyaluronic acid, collagen, fibrin, whey protein, gelatine, and sodium alginate [19-21]. Collagen and hyaluronic acid are both found in the extracellular matrix (ECM) within the human body and have been used in tissue engineering for a range of purposes, such as endothelial and liver cell 3D culture and cornea regeneration [22-25]. However, these ECM polymers are sourced from animals, have issues with printability and are often soft [26, 27]. Sodium alginate is a more readily available ionic polymer derived from brown seaweed [28]. Due to its high availability, purity, printability, and biocompatibility, it is often used as a bioink in bioprinting applications [28, 29]. Whilst alginate has been shown to form a hydrogel at a range of concentrations it can only be easily bioprinted at low concentrations (5-15%) due to an increase in viscosity [28, 30].

One potential route to creating stiffer printable materials is to use the non-Newtonian flow behaviour that many polymeric materials possess. Shear-thinning hydrogels display reduced viscosity in response to shear strain which allows them to be printed without damaging encapsulated cells (Figure 1) [31-35]. Whilst many natural polymers display some shear-thinning properties, this effect is often subtle and reliant on the low viscosity of the hydrogel to protect encapsulated cells during printing [21]. Nanofibrillar materials on the other hand have been shown to possess more dramatic shear-thinning properties, likely due to fibrillar disentanglement or the alignment of fibres upon the application of shear force [36]. For this reason, we have investigated adding nanofibrillar assemblies formed from self-assembled peptides (SAPs) to form hybrid bioinks. SAPs typically consist of 2-20 amino acids that undergo spontaneous assembly into ordered nanostructures such as nanotubes, fibres, sheets, and vesicles [37-39]. SAPs make attractive biomimetic scaffolds due to their ability to assemble into nanofibrils resembling the ECM [38, 40-42]. They are often tolerant to the introduction of additional chemistries such as cell recognition sequences [41, 43]; they have highly attractive self-healing properties [44-46] and their synthetic nature means they can be prepared in high purity [47]. However, a major limitation with SAP hydrogels is their non-covalent cross-linking, meaning they are generally very soft and susceptible to shear forces, such as those generated during bioprinting, which can break the hydrogel by disrupting the non-covalent bonds holding the gel together [48].

**Figure 1.**
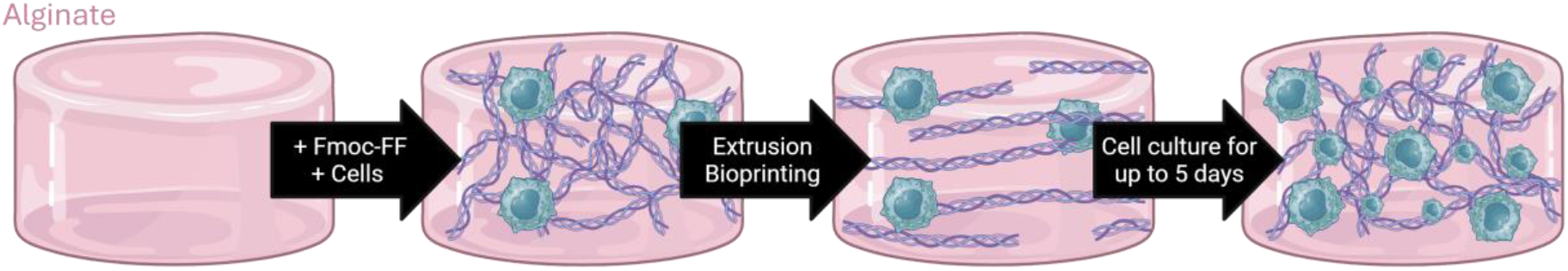
The formation of macrophage laden hybrid alginate and Fmoc-FF hydrogels.

To address the respective limitations of natural polymers and SAPs, hybrids can be developed to create an optimised material. Various components can be incorporated into the hydrogels to change its chemical and mechanical nature. To date there is only a few examples of hybrid materials composed of SAPs and natural polymers finding applications in bioprinting thus this is an important area for further exploration [49-52].

In this work, we have developed hybrid bioinks formed from the SAP Fluorenyl-9-methoxycarbonyl (Fmoc)-diphenylalanine (Fmoc-FF) [39, 53-59] and the natural polymer sodium alginate. The addition of small amounts (0.5 % w/v) of Fmoc-FF forms hybrid bioinks that after covalent crosslinking of the sodium alginate possess stiffnesses around 8 times that of alginate alone. These hydrogels show much improved printability, support the growth of encapsulated macrophage cells over 5 days, and have exciting self-healing properties. Further, we provide important insights into how the organic solvent used to initially solubilise the Fmoc-FF has significant effects on both the biocompatibility and the physiochemical properties of the hybrid bioinks. The development of these new stiff, tuneable, and printable SAP-bioinks has applications in tissue engineering, biosensors, wearable biomaterials, wound healing gels and bioengineering.

## 3. Experimental (Methods and materials)

### 3.1 Hydrogel preparation

Alginate hydrogels were formed by dissolving sodium alginate powder (Sigma-Aldrich) in ultra-pure water (18.2 MΩ·cm) (Milli-Q®) water at 5-20 % (w/v) concentrations. The samples were left on a rolling machine to mix for 5-60 minutes depending on the concentration. Samples were crosslinked with 50 mM calcium chloride (CaCl_2_) (Chem Supply) for 5-60 minutes and then stored in tris buffered saline (TBS) (Sigma-Aldrich).

Fmoc-FF hydrogels were formed by solubilising Fmoc-FF peptides (Bachem, Switzerland) with dimethyl sulfoxide (DMSO) or 1,1,1,3,3,3-hexafluoroisopopanol (HFIP) (Sigma-Aldrich) to form a 100 mg/ml stock solution. Solubilised peptides were diluted to 5 mg/ml in ultra-pure water (18.2 MΩ·cm) (Milli-Q®) and vortexed for 20 seconds, and then left to equilibrate overnight.

To form the hybrid bioinks, the Fmoc-FF stock solution was diluted to 0.5% (w/v) in a 5% (w/v) uncrosslinked alginate solution, and vortexed for 20 seconds. The samples were left to equilibrate overnight. Following this, samples were crosslinked for 5-60 minutes using 50 mM CaCl_2_ (Chem Supply) and then stored in TBS (Sigma Aldrich).

### 3.2 Rheological measurements

To assess the physical properties of the crosslinked scaffolds, amplitude sweeps, 8 interval thixotropic tests (8-ITT) and viscosity tests were performed using a parallel plate rheometer (Anton Paar MCR302). Solid (crosslinked) samples were analysed using a 10 mm parallel plate (PP-10) and fluid (uncrosslinked) samples were analysed using a 25 mm 1° cone plate (CP1-25). Amplitude sweeps were designed with 21 data points and performed from 0.01-100% shear strain, with a set angular frequency of 10 rads^-1^ and a force-controlled moving profile set to 0.1 N of constant applied force. Using a force-controlled profile rather than a set gap ensured consistency between samples regardless of subtle height differences. 8-ITT were performed using 8 cycles alternating from high strain (100 %) to low strain (0.1 %) with 4 seconds per point and 10 points per cycle. The tests were also performed with a force-controlled moving profile set to 0.1 N of constant applied force. The hydrogels viscosity was measured using a 25 mm 1° cone plate (CP1-25). The viscosity curves were designed with 30 data points, a shear rate range of 0.001 to 1000 (1/s) and a set angular frequency of 10 rads^-1^. All tests were performed at 21° C.

### 3.3 3D Bioprinting

Hydrogels were loaded into pneumatic print cartridges (Cellink) using female-female luer adapters (Cellink) and placed into the BIOX 3D bioprinter (Cellink) for printing. Dispensing nozzles (Cellink) from 20-25 gauge were used depending on the hydrogel. Extrusion pressure was between 10-200 kPa, print height was 0.1 mm from plate, and print speed varied depending on the biomaterial and shapes being printed (1-5 mm/s). All printing was performed at 21° C.

### 3.4 Thioflavin T imaging

Thioflavin T (ThT) dye (Sigma Aldrich) was used to visualise Fmoc-FF fibres within the hybrid scaffolds with laser scanning confocal microscopy. 200 µL samples were bioprinted into 8 chamber microscopy slides (Lab-Tek) and then stained overnight with 25 µM ThT and washed 6 times with TBS before being imaged. Zeiss LSM 780 microscope (Zeiss, Germany) was used to image the samples at an excitation wavelength of 450 nm and an emission wavelength of 485 nm and a 40x water immersion lens (NA = 1.1) at room temperature.

### 3.5 Scanning Electron Microscopy

Prior to scanning electron microscopy (SEM) imaging, 2 mm hydrogel droplets were bioprinted and crosslinked for 5 minutes with 50 mM CaCl_2_. The samples were then washed for 10 minutes with 50%, 60%, 70%, 80%, 90% and 100% ethanol before being dried in a CO_2_ E3100 critical point dryer (Quorum Technologies) for 30 minutes. The dried samples were mounted to SEM mounts using conductive double-sided tape and platinum sputter coated to a thickness of 5 nm. The samples were then imaged in the Hitachi FE-SEM SU7000 (Hitachi) using 3-5 kV at 1000-20,000 K magnification.

### 3.6 Transmission Electron Microscopy

Copper 200 mesh square grids with a formvar-carbon support film (GSCU300CC-50, ProSciTech, Qld, Australia) were glow discharged for 60 s in an Emitech k950x with k350 attachment. Hydrogel samples (∼ 5 μL) were placed onto the grids and with the excess blotted off to create a thin film. Samples were crosslinked briefly with 50 mM CaCl_2_. Two drops of uranium acetate (2 %) was then used to negatively stain the hydrogel samples, with excess uranium acetate removed by blotting with filter paper after 10s. Grids were imaged using a Joel JEM-2100 (JEOL, Australasia) Pty Ltd) transmission electron microscope equipped with an AMT Nanosprint15 Mk-II (Newspec Pty Ltd. Adelaide, Australia).

### 3.7 Thioflavin T fluorescence plate reader assay

Thioflavin T (ThT) dye (Sigma Aldrich) was used to measure the presence of Fmoc-FF fibres in hybrid scaffolds over time. 100 µL samples of hybrid hydrogels were pipetted into 96 well plates and crosslinked for 5 minutes with 50 mM CaCl_2_. On days 0, 1, 3 and 5 25 µM ThT dye was added and left at 4 °C for 6 hours, then rinsed four times with TBS. The fluorescence spectra was then recorded using a CLARIOstar plate reader (BMG Labtech) (excitation 435 nm, emission 475 nm). Alginate-only samples were tested and showed minimal fluorescence, graphed data have had alginate-only values subtracted.

### 3.8 Cell viability assays

#### Cells seeded on top of hydrogels (2D cell culture)

Hybrid scaffolds were bioprinted into 96 well plates and crosslinked using 50 mM CaCl_2_. Scaffolds were washed six times with DMEM (Gibco), supplemented with 1% penicillin and streptomycin (Gibco), 10% foetal bovine serum (FBS) (Gibco) and 2 mM glutamine (Gibco) before leaving in the media overnight. Murine RAW264.7 cells were then seeded on the surface of the scaffolds at 250,000 cells/mL. Resazurin-based AlamarBlue-HS (Sigma-Aldrich) dye was added 2 hours before the plate read and incubated at 37 °C 10 % CO_2_. Post incubation fluorescence spectra was recorded on days 0, 1, 3 and 5 using a CLARIOstar plate reader (BMG Labtech) (excitation 560 nm, emission 590 nm) at 25 °C using the top optic. Scaffold-only controls were used to subtract background fluorescence readings from samples.

#### Cells within hydrogel (3D cell culture)

Hydrogels were made as per the hydrogel preparation methods above but with DMEM (Gibco) supplemented with 1 % penicillin and streptomycin (Gibco), 10 % FBS and 2 mM Glutamine (Gibco) instead of ultra-pure water. They were then gently mixed with a stirring flee for 1 hour on heat block at 40 °C before being placed into bioprinting syringe. Murine RAW264.7 macrophages were added to the hydrogels at a concentration of 1 million cells/mL and left in incubator at 37 °C 10 % CO_2_ for 2 hours. The newly formed bioinks were then bioprinted into 96 well plates and crosslinked using 50 mM CaCl_2_. Bioinks were washed 3 times with the same supplemented DMEM they were made with above. The bioinks were left to rest for 4 hours. Resazurin-based AlamarBlue-HS dye was added 2 hours before the plate read and incubated at 37°C 10% CO_2_. Post incubation fluorescence spectra was recorded on days 0, 1, 3 and 5 using a CLARIOstar plate reader (excitation 560 nm, emission 590 nm) at 25°C using the top optic. Scaffold-only controls were used to subtract background fluorescence readings from samples.

### 3.9 Fluorescence imaging of encapsulated cells

RAW264.7 macrophages were stained with 10 mM CellTracker Orange CMRA (Thermo Fisher) for 45 minutes before being incorporated into alginate and hybrid hydrogels (made as per the 3D cell viability assay hydrogel formulation steps) at a concentration of 4 million cells/mL to form cell-laden bioinks. 200 µL samples were bioprinted into 8 chamber microscopy slides (Lab-Tek) and crosslinked with 50 mM calcium chloride for 5 minutes and then imaged with a Zeiss LSM 780 microscope (Zeiss, Germany) at an excitation wavelength of 548 nm and an emission wavelength of 576 nm with a 40x water immersion lens (NA = 1.1) at room temperature (21°C). Imaris software (Imaris, Oxford Instruments) was used to create 3D representations of the z-stacks.

### 3.10 Statistics

Statistical analysis was performed in GraphPad Prism using one and two-way ANOVA (with Tukey comparison). Each experiment was independently performed 3 times, with 3 technical replicates per experiment.

## 4. Results and discussion

### 4.1 The addition of Fmoc-FF to alginate hydrogels increases their stiffness whilst simultaneously improving their printability

To investigate if we could improve the printability of Fmoc-FF containing hydrogels we formed hybrid materials where the aqueous phase of the gel was replaced by a viscous solution of the natural biopolymer sodium alginate. In addition, as it is known that the organic solvent used to initially dissolve the Fmoc-FF peptides can have a significant effect on the resultant assemblies [39, 58], we compared hybrid hydrogels that were made from stock peptide solutions dissolved in either DMSO or HFIP (hereon referred to as DMSO or HFIP hybrids respectively).

By varying the relative concentrations of alginate and Fmoc-FF, we identified that 5 % (w/v) alginate solution in water mixed with 0.5 % (w/v) Fmoc-FF (for both DMSO and HFIP hybrids) produced hydrogels that maintained their 3D structure over multiple hours (Figure S1). Using parallel plate rheology, we compared the storage modulus (G’) in the linear viscoelastic region (LVR) of the hybrid hydrogels to that of the alginate alone (Figure S2). Before crosslinking of the alginate component of the hybrid hydrogels, the addition of 0.5 % Fmoc-FF increased the G’ by a factor of 277 (figure 2a) when compared to 5% alginate alone, and there was no significant difference between the G’ of the DMSO or HFIP hybrids. Even after increasing the alginate concentration to 20 % the G’ of the alginate gels remained lower than the hybrids, which contain only 5 % alginate (Figure 2a). After crosslinking this effect was more pronounced in the DMSO hybrids, where the addition of Fmoc-FF increased G’ by almost 10 times compared to 5 % alginate alone and was almost double that of 20 % alginate hydrogels (figure 2b). The increase in storage modulus was not as pronounced in the HFIP hybrids, however these gels still possessed a G’ of 25 kPa, 8-times stiffer than the alginate gels alone, at 5 % (Figure 2b). This shows that the addition of relatively low concentrations of SAPs can significantly increase the stiffness of natural polymer hydrogels, allowing the fabrication of materials with stiffnesses approaching 40 kPa.

**Figure 2:**
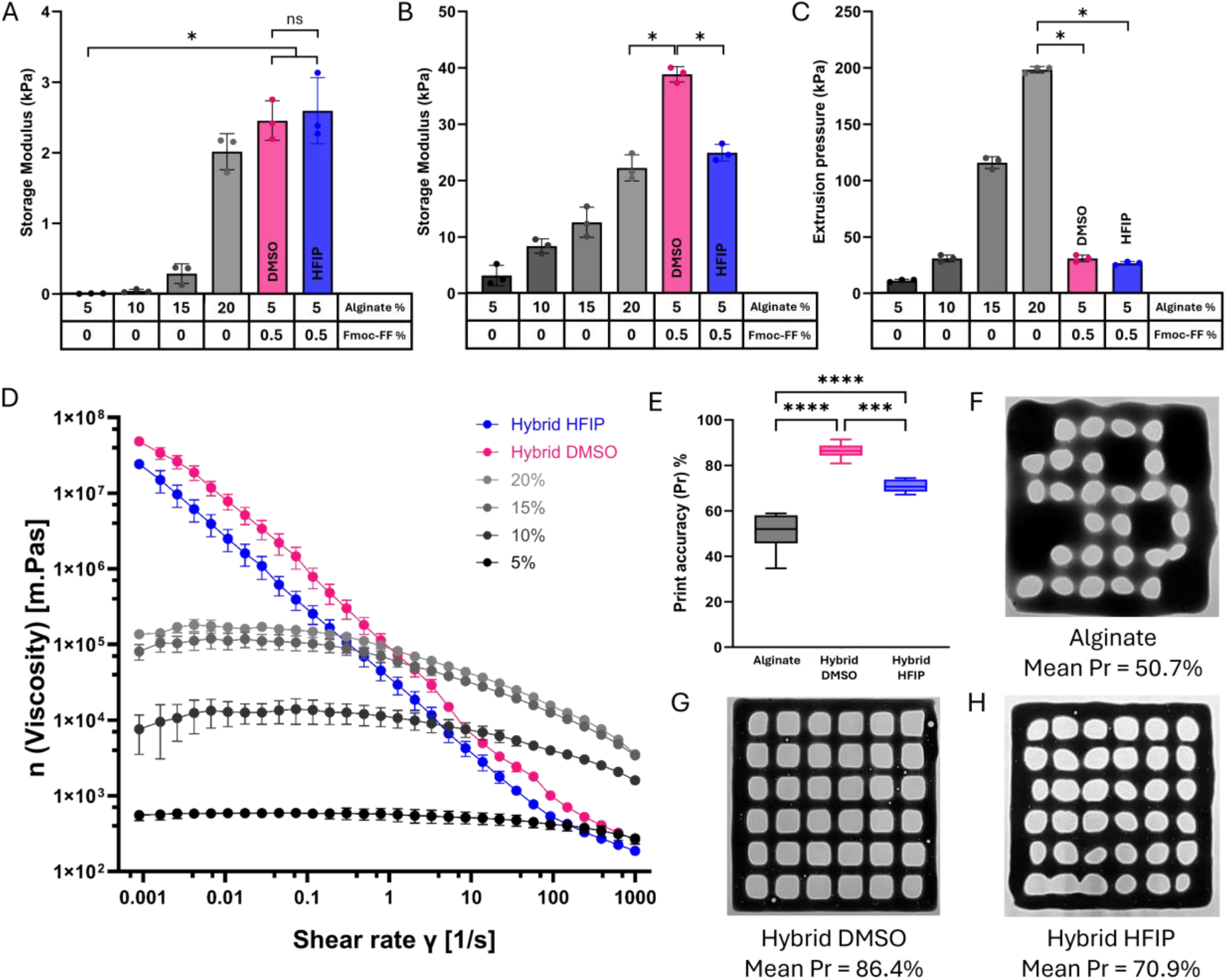
The addition of Fmoc-FF peptides to alginate hydrogels increases hydrogels stiffness before and after crosslinking whilst also improving printability. The storage modulus of alginate and hybrid hydrogels at 0.1 % shear strain before crosslinking (a), after crosslinking (b), and the extrusion pressure required to print the hydrogels (c). Viscosity curves of alginate and hybrid hydrogels (d). Print accuracy values of alginate, hybrid DMSO and hybrid HFIP printed mesh constructs (as determined from equation 1) (e). Representative images and mean print accuracy values of 3D printed hydrogel mesh constructs (f-h). Bars and data points represent the mean of 3 independent biological replicates (n=3) and error bars show the standard deviation of the mean. * indicates p-value <0.05, *** <0.001, ****<0.0001.

Whilst the ability to produce hydrogels with a variety of stiffnesses is important in regenerative medicine and 3D cell culture, it is also highly desirable that these materials remain printable at extrusion pressures that are compatible with the bioprinting of cell-laden hydrogels (bioinks) [60]. Therefore, we investigated the minimum extrusion pressure required to print a constant stream of our hybrid gels (figure 2c and S1). Alginate gels alone required a large increase in extrusion pressure as the concentration of alginate increased (figure 2c). At 20 % alginate almost 200 kPa of pressure was required; at this pressure, encapsulated cells within a cell-laden bioink would experience cytotoxic shear forces [61]. By contrast, the addition of 0.5 % Fmoc-FF had a remarkable effect on the extrusion pressure required to print the hybrid gels. Compared to hydrogels with equivalent high stiffness, such as 20 % alginate, the 0.5 % Fmoc-FF hybrids required 6 times less extrusion pressure to be printed (figure 2c).

To investigate the origin of the increased printability of the hybrid hydrogels we compared the rate of change of viscosity with increasing shear rate between the two hybrids and 5 % alginate (figure 2d). For 5 % alginate gels alone there was very little change in viscosity upon increased shear, indicating it is a Newtonian fluid. For both the DMSO and HFIP hybrids, we observed a large drop in viscosity with increasing shear rate, indicating that both bioinks are highly shear-thinning. The shear-thinning behaviour of the hybrid gels explains their much-improved printability compared to the alginate scaffolds, suggesting that these materials may have important applications where it is desirable to print materials with a range of mechanical properties. This includes not just tissue engineering and regenerative medicine, but also fields like biosensors and wearable technologies.

To examine the macroscale architecture of the printed constructs we imaged grid meshes to compare the print fidelity of alginate versus the HFIP and DMSO hybrid hydrogels (Figure 2e-h). The morphology of the interconnected channels was analysed using Fiji software to determine the total area of the pores, which was then used in the following equation to calculate the % print accuracy (𝑃𝑟) of the different materials:

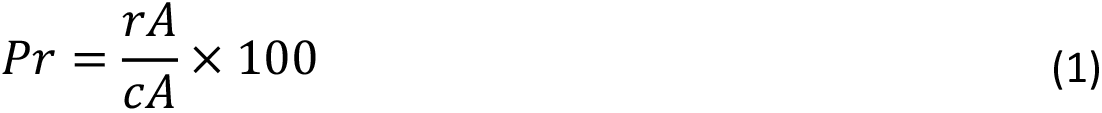

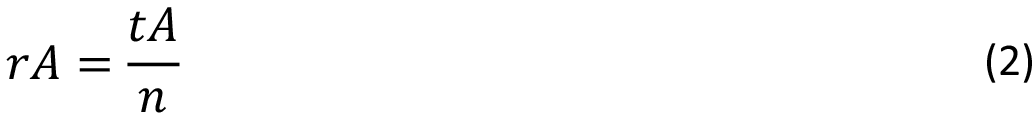

Where 𝑟𝐴 is the relative individual pore area, 𝑐𝐴 is the calculated individual pore area, 𝑡𝐴 is the total pore area and 𝑛 is the proposed number of pores.

The DMSO hybrids had the highest print accuracy (86.4 %), followed by the HFIP hybrids (70.9%). Whereas, the 5 % alginate hydrogels possessed a much lower print accuracy of 50.7 % (figure 2e), indicating that the incorporation of Fmoc-FF peptides into the hybrid materials improves their printing quality while still retaining low extrusion pressures required for cellular applications.

### 4.2 The hybrid gels have self-healing properties

Shear-thinning materials can also have self-healing properties that are highly attractive in tissue engineering, regenerative medicine, and wound dressing applications [45, 62, 63]. To investigate the self-healing potential of our hybrid materials, we performed rheological thixotropic interval tests where the gels were subjected to cycles of low and high strain (Figure 3 a-c). In the high strain regions, the microstructure of the hydrogels is broken, resulting in a phase transition from a viscoelastic solid (G’>G”) to a viscoelastic liquid (G”>G’). However, if the material is self-healing, when the high strain is removed, we expect to see a rapid return to a viscoelastic solid and the maintenance of a significant gap between G’ and G” (Table 1). In the case of the alginate-only hydrogels we observed a loss of self-healing ability where after 4 low strain cycles the material did not reform to a viscoelastic solid, instead remaining a liquid (G’’>G’) (Figure 3a). By contrast, the DMSO hybrids exhibited negligible change in G’ across the four cycles of low strain, and there was minimal change in the gap between G’ and G’’ across the cycles (Figure 3b, table 1). For the HFIP hybrids, this self-healing property, whilst still apparent, was less pronounced with a 58 % drop in G’ after 4 low strain cycles and a significant narrowing of the gap between G’ and G’’ (37 % decline) (Figure 3c, table 1). In summary, the addition of SAPs increased the self-healing properties of alginate hydrogels, likely due to the thixotropic nature of the self-assembled Fmoc-FF nanofibrils [64]. For reasons investigated in the remainder of this manuscript this self-healing property was improved in the DMSO hybrids compared to the HFIP hybrids.

**Figure 3:**
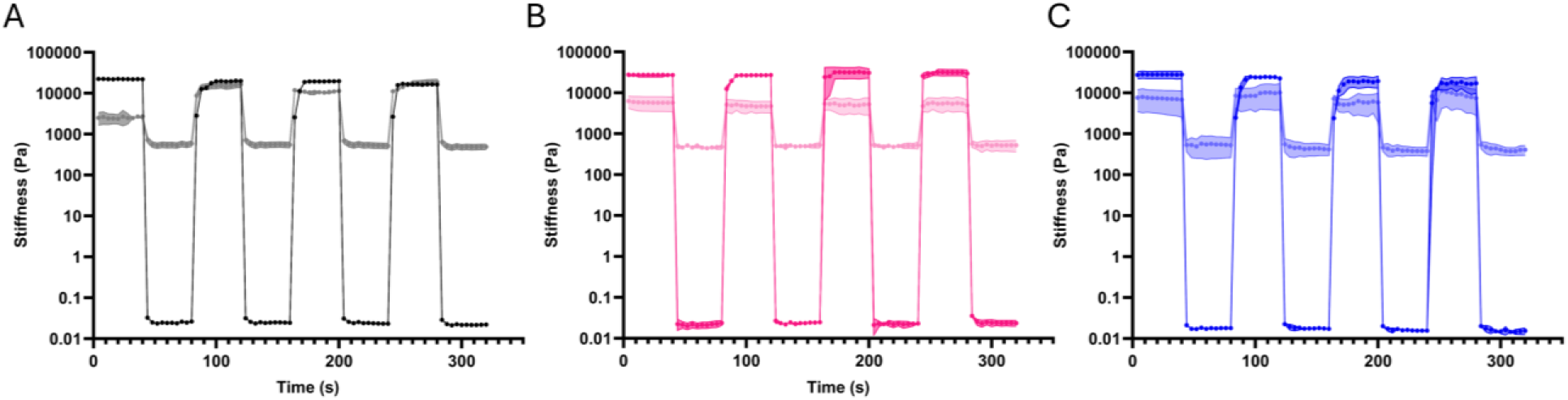
Hybrid hydrogels have improved self-healing properties. 8-interval thixotropic tests of crosslinked alginate (a), DMSO hybrids (b) and HFIP hybrids (c) at alternating high (100%) and low (0.1%) strain. Data points represent the mean of 3 independent biological replicates (n=3) and error bars show the standard deviation of the mean.

**Table 1.**
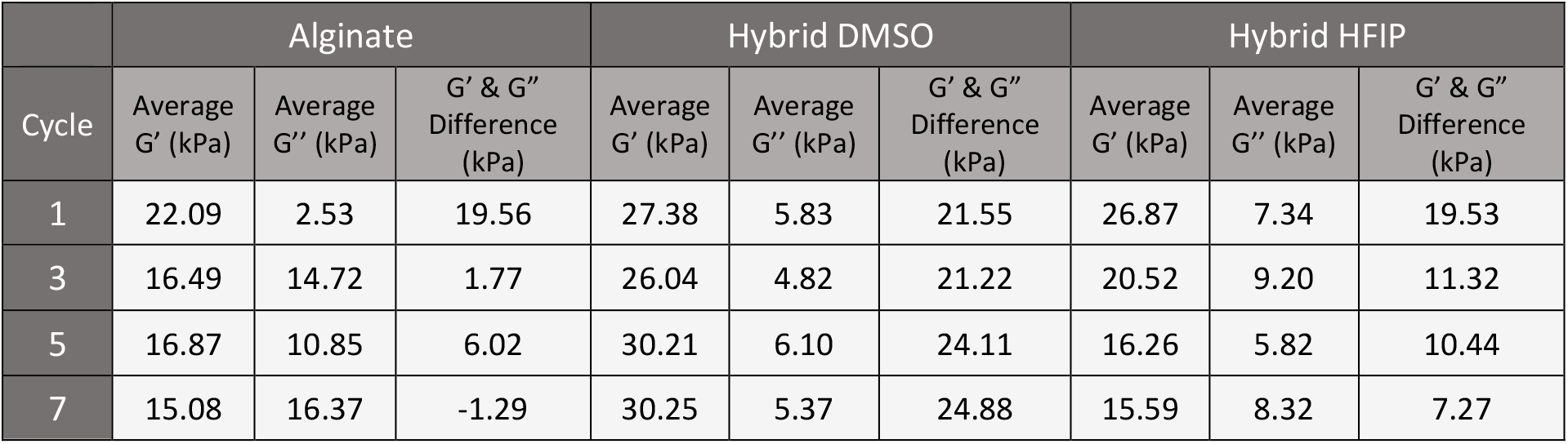
Average storage modulus (G’) and loss modulus (G’’) of alginate, DMSO hybrids and HFIP hybrids at low strain (0.1%) during 8-interval thixotropic tests.

### 4.3 The hybrid bioinks support the growth of 3D printed encapsulated macrophages

In this proof-of-principle work, we chose a macrophage cell line as our choice of encapsulated cells. Macrophages were chosen as, similarly to much more well-studied mesenchymal stem cells, they are likely to be very sensitive to their 3D environment and its mechanical properties [65, 66]. As tissue resident macrophages are present all throughout the body in tissues with varying stiffness it is highly likely that the mechanical properties of those tissues play a vital role in directing their function.[65] Therefore, to ascertain if the hybrid bioinks supported the printing and subsequent growth of macrophages we imaged RAW264.7 cells within the hydrogels and on glass after 24 hours (figure 4a-b, figure S3). Cells were visible in both HFIP and DMSO hybrids which had a rounded morphology (cf. the more spread cells on glass in figure S3) typical of macrophage cells cultured in 3D (figure 4a-b) [66]. To quantify long-term cell viability, we formed cell-laden bioinks with RAW264.7 macrophages and measured their viability over 5 days (termed 3D culture, figure 4c) compared to the cell growth of the same cell line seeded on top of the hybrid bioinks (termed 2D culture, figure S4). Despite a small amount of cell growth after 1 day, 3D cultured cells in the DMSO hybrids had a decreased viability at all further time points, which was consistent with 2D cultured cells (figure S4). We hypothesised that this was due to residual DMSO becoming trapped within the scaffold due to the increased viscosity of the hybrid materials, as such toxicity generally isn’t observed in Fmoc-FF-only gels [67]. When the same experiments were performed with HFIP hybrids, cells cultured in 2D and 3D were viable over the time course, with only a small drop in viability at 5 days likely due to the cells using all available nutrients. The improved viability of cells cultured in the HFIP hybrid scaffolds compared to alginate alone is perhaps somewhat surprising as no specific chemistries have been added with the SAP assemblies to promote cell adhesion and viability. Instead, we suggest that the nanofibrillar morphology of these structures is promoting cell growth through improved biomimetic ECM mimicking morphologies of the SAP in the hybrid bioink [68]. The improved ability of the HFIP scaffolds to encapsulate cells and act as biocompatible bioinks (cf. the DMSO hybrids) is likely to be due to the increased volatility of HFIP compared to DMSO. To confirm that the source of the cell toxicity was residual organic solvent trapped within the materials, we cultured the same cell line on alginate only gels doped with the same concentration of DMSO or HFIP (5 % v/v) and observed a similar decrease in cell viability in cells seeded on the DMSO hybrid materials, but not the HFIP hybrids (Figure S5). Overall, the cell viability and microscopy show that these hybrid bioinks support the growth of macrophages after their extrusion through a printing head.

**Figure 4:**
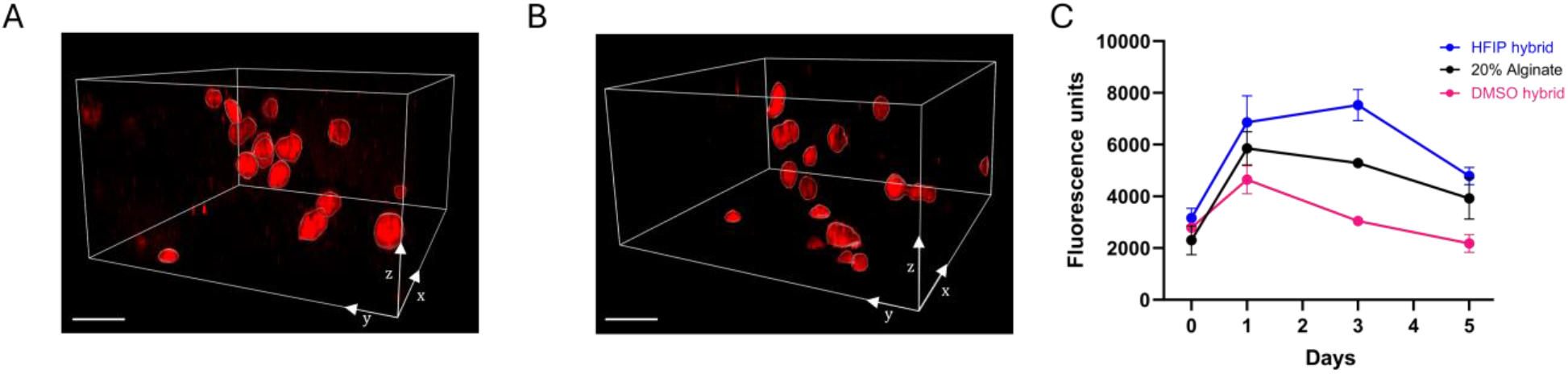
HFIP hybrid hydrogels support long-term cell viability. Laser scanning confocal microscopy images of RAW264.7 cells stained with CellTracker Orange inside DMSO hybrid (a) and HFIP hybrid (b) hydrogels after 24 hours. Viability of RAW264.7 macrophages within cell-laden 3D printed hydrogel scaffolds over 5 days (c). Data points represent the mean of 3 independent biological replicates (n=3) and error bars show the standard deviation of the mean. Scale bars represent 40 µm.

### 4.4 The choice of organic solvent has significant effects on the architecture of the printed hydrogels

We next examined the microarchitecture of the HFIP and DMSO hybrid hydrogels and performed scanning electron microscopy (SEM) to examine their surface morphology. A range of topographical differences were observed across the surface of all three materials (figure 5a-c). DMSO hybrid scaffolds (figure 5a) had a more fibrillar appearance when compared to alginate alone (figure 5b), which had a highly corrugated surface. By contrast, the HFIP hybrids (figure 5c) had multiple holes or tears in their surface (pink arrows, figure 5) which was unlike both the alginate and DMSO hybrids. We postulate that these tears were formed due to Marangoni forces generated by the HFIP escaping the hydrogel, similar to observed in Pena-Francesch *et. al.[69]* HFIP is a highly volatile solvent with very low surface tension, thus when alginate is crosslinked residual HFIP is forced to the surface creating large gradients in interfacial tension between the surface of the materials and the aqueous environment, leading to the tears observed in the SEM images. This forceful expulsion of HFIP from the hybrids likely explains why there was no decrease in macrophage viability in the same way as the less volatile DMSO.

**Figure 5:**
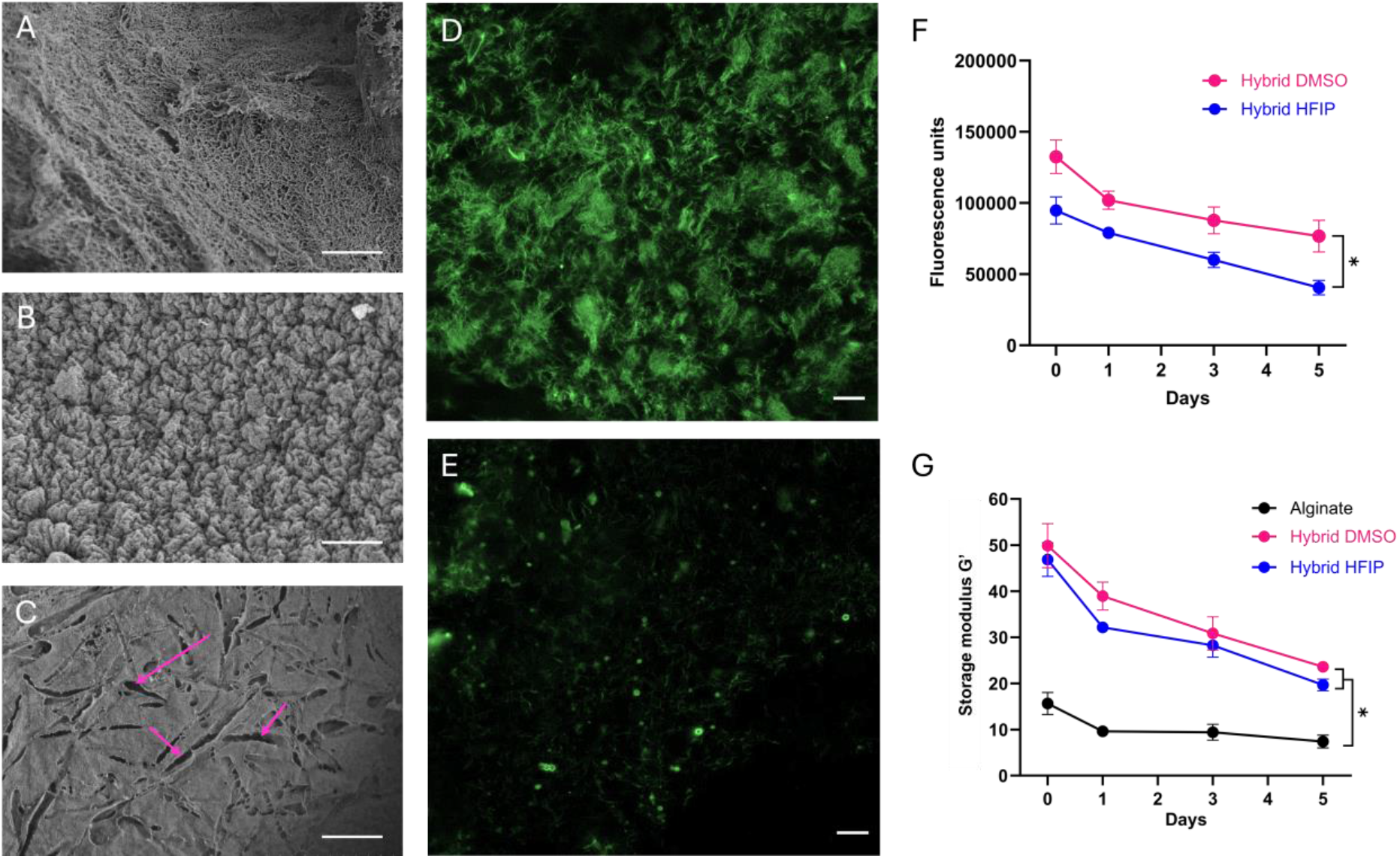
Microscopic differences emerge between DMSO and HFIP hybrid hydrogels. Scanning electron microscopy (SEM) of critical point dried DMSO hybrids (a), alginate (b) and HFIP hybrids (c). LSCM images of crosslinked DMSO hybrids (d) and HFIP hybrids (e). Thioflavin T fluorescence of Fmoc-FF fibres within HFIP and DMSO hybrids over 5 days (f). Degradation of alginate and hybrid hydrogels over 5 days (g). Scale bars represent 2 µm (a-c) and 25 µm (d-e). Data points represent the mean of 3 independent biological replicates (n=3) and error bars show the standard deviation of the mean. * indicates p-value <0.05 The data above highlighted significant differences between the interior and surface of the hybrid materials made with Fmoc-FF solubilised in HFIP and DMSO. ThT data showed that the HFIP hybrids formed a less extensive network of Fmoc-FF fibrils (figure 5e-f) resulting in reduced thixotropic self-healing ability (figure 3b), and a faster degradation of mechanical properties (figure 5g). Additionally, due to the Marangoni forces generated by escaping HFIP, we observed tears in the surface of the HFIP hybrids (figure 5c). The remainder of this study is dedicated to characterising and explaining the apparent reduction of ThT positive nanofibrillar assemblies when Fmoc-FF scaffolds are prepared using HFIP.

We next investigated the internal microstructure of the hydrogels by laser scanning confocal microscopy (LSCM) and Thioflavin T (ThT) staining. ThT binds to the extended β-sheet structures in the Fmoc-FF nanofibrils, becoming highly fluorescent [70]. The levels and distribution of ThT fluorescence between the DMSO and HFIP hybrids varied significantly (figure 5d-e), where the HFIP hybrids had a much sparser distribution of ThT positive regions (figure 5e). We quantified the amount of ThT fluorescence in both hybrids over time with a fluorescent plate reader (figure 5f). Both hybrid materials show a similar decrease in ThT fluorescence over 5 days, however the DMSO hybrids had a significantly greater fluorescence at all 4 time points. This data combined with the fluorescence microscopy suggests that β-sheet rich Fmoc-FF fibrils form to a greater extent in the hybrid DMSO scaffolds compared to the hybrid HFIP scaffolds.

The mechanical stability of the hydrogels influences the materials potential future applications. Therefore, we measured the change in the G’ of crosslinked scaffolds over 5 days (Figure 5g). The alginate-alone and the hybrid hydrogels had a similar amount of degradation, decreasing G’ by 47 % and 42 % over 5 days, respectively. Whilst only the DMSO hybrids maintained a stiffness above 20 kPa at day 5, both hybrid materials had a significantly greater stiffness than alginate alone at all timepoints. This decrease in mechanical stability over time correlates with the reduced ThT fluorescence, suggesting that the Fmoc-FF fibrils disassemble over a few days. Different rates of degradation may be advantageous depending on the application of the materials. For example, it is desirable with implantable sacrificial scaffolds to match the degradation rate of the scaffold to new tissue growth [71]. By contrast, for long-term implants or drug delivery models, a more stable scaffold may be more applicable [72-74].

To determine the mechanical tuneability of the hydrogels, we varied the ionic crosslinking times to determine its effect on the stiffness of hydrogels. We observed a positive correlation between crosslinking time and G’ for all materials (Figure S7). The G’ of alginate hydrogels increased by a factor of 7.3 over 60 minutes whereas the DMSO and HFIP hybrids increased by a factor of 8.7 and 3.8, respectively. This data shows that the mechanical properties of both bioinks can be tuned over a physiologically relevant range of tissue stiffnesses (∼ 3-65 kPa), which may be desirable in creating bioinks that mimic a range of clinically important tissues, such as breast, endothelial, muscle and cartilage tissue [75].

### 4.5 Solvent specific non-covalent interactions promote crystallisation via an Ostwald ripening effect in the HFIP scaffolds

To investigate if the HFIP hybrids possessed fewer β-sheet rich nanofibrils, we analysed the hybrid materials and the Fmoc-FF assemblies alone by transmission electron microscopy (TEM) (figure 6a-h). DMSO hybrids at 24 hours had an intricate matrix of alginate polymer chains (figure 6a-d, purple arrows, also observed in alginate only samples, figure S8) and large Fmoc-FF fibres in multi-fibrillar ribbons (figure 5a-d, cyan arrows); similar in morphology to previously reported Fmoc-FF fibrils [54, 55, 76]. The HFIP hybrids had a reduction in the number of Fmoc-FF fibrils at 24 hours, with an increase in the amount of large crystal-like Fmoc-FF structures (figure 6d, green arrows). This reduction in the amount of fibres explains the reduced ThT fluorescence (figure 5g).

**Figure 6:**
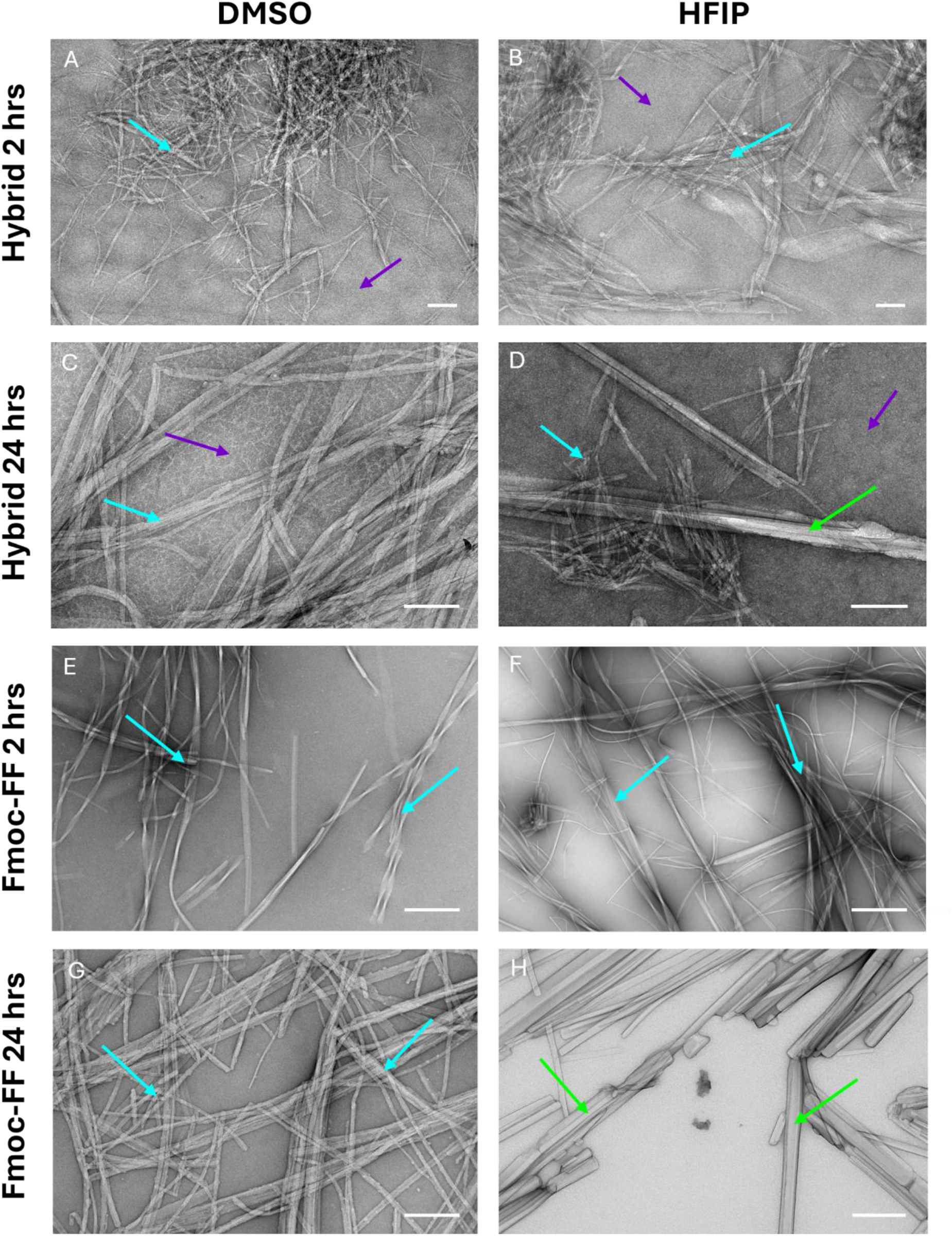
Crystalline structures are present in HFIP hybrids and Fmoc-FF only gels when made using HFIP as a peptide solvent. Transmission electron microscopy images of HFIP and DMSO hybrid hydrogels and Fmoc-FF only hydrogels made with DMSO and HFIP at 2 and 24 hours post gelation. Cyan arrows show Fmoc-FF fibres, purple arrows show alginate polymers and green arrows show Fmoc-FF crystalline structures. Scale bars represent 200 nm.

The above results show that the morphology of the Fmoc-FF assemblies varies depending on the choice of initial solvent to solubilise this peptide. It is known that the Fmoc group in Fmoc-FF is labile and can easily dissociate in the presence of, for instance, high pH [77]. Both Fmoc-FF and FF alone have been reported to form a highly polymorphic array of nanostructures including fibrils, crystals, and amorphous aggregates [39, 53, 78]. As HFIP is an unusually reactive solvent we performed mass spectrometry experiments to determine if the Fmoc group of monomeric Fmoc-FF was cleaved when the peptide is dissolved in HFIP, as this would explain the higher tendency for crystal formation for materials made from HFIP stocks (figure S9) [79] [80]. However, the mass spectra showed no evidence of increased Fmoc dissociation in HFIP solutions of Fmoc-FF when compared to the peptide dissolved in DMSO, therefore is unlikely to explain the significantly different structures formed.

To further investigate the differences in the polymorphic assembly of the DMSO and HFIP hybrids we compared their morphologies after different assembly times. At 2 hours both the DMSO and HFIP hybrids possessed similar mixtures of nanofibers and multifilamentous ribbons (figure 6a-b, cyan arrows) alongside alginate polymers (figure 6a-b, purple arrows). It is only after 24 hours that the crystalline polymorph is evident in the HFIP hybrids. To confirm that this conversion from a fibrillar to a crystalline polymorph was not due to the presence of alginate in the hybrid scaffolds, we analysed Fmoc-FF only gels made with DMSO and HFIP by TEM after 2- and 24-hours assembly. Similar to the hybrid scaffolds, we observed primarily fibrillar assemblies at 2 hours, and at 24 hours crystalline assemblies were only present in the hydrogels formed from HFIP stocks (figure 6h, green arrows).

Based on these findings, we suggest that the formation of crystals within the HFIP hybrids is time-dependent and follows an Ostwald ripening process, where small fibres are initially formed (2 hours) but then are replaced by large crystal structures which are more thermodynamically stable (> 24 hours). Dudukovic *et al* identified this process and found that different solvents altered the phase behaviour of Fmoc-FF peptides and intramolecular non-covalent bonding [39]. They noted that crystals formed when there were higher levels of hydrogen bonding and dispersion forces but fewer polar bonds; as we observe with HFIP materials. The presence of crystals and decrease in fibres in the HFIP hybrids also helps explain the decrease in the physical stability of the biomaterials (figures 2-5), as the fibres provide the hydrogels’ elasticity, stability, and self-healing properties [81, 82]. Ultimately, we found the use of HFIP as a solvent in hybrid hydrogels, whilst improving their biocompatibility, had pronounced effects on the physical and molecular properties of the resultant material due to the formation of crystalline Fmoc-FF structures instead of fibres.

## 5. Conclusions

By combining the self-assembled peptide Fmoc-FF and sodium alginate we have developed a biocompatible, self-healing, printable hybrid biomaterial with tuneable mechanical properties. The addition of 0.5 % self-assembled peptide permits the bioprinting of stiff hydrogels at printing pressures 5 times less than what is required for alginate only gels of equivalent stiffness. This greatly improves the ability to extrude cell-laden bioinks without inducing a toxic response. These highly printable, self-healing, biocompatible bioinks may have applications in smart wound dressings, injectable hydrogels for drug delivery, and for understanding the behaviour of cells (particularly macrophages) in tissue mimetic 3D *in vitro* environments. We also show that the choice of organic solvent used to dissolve the self-assembling peptides can significantly impact not only the nanoarchitecture of the assemblies, but also the physical properties of the resulting biomaterials, highlighting the importance of the careful choice of solvent in the fabrication of biomaterials.

## Supporting information

Supplemental Information for Field et al

## Acknowledgements

We would like to acknowledge Rohan Lowe from the Latrobe University Proteomics and Metabolomics Platform for the assistance with mass spectrometry. This research is supported by an Australian Government Research Training Program (RTP) Scholarship. Dr Reynolds would like to thank the La Trobe Institute for Molecular Sciences (LIMS) for a fellowship that financially supports this work. We acknowledge figure 1, and the table of contents image was created with BioRender.com

