## Supplemental Information for Field et al for "Self-healing, biocompatible bioinks from self-assembled peptide and alginate hybrid hydrogels"

1. Department of Biochemistry & Chemistry, La Trobe Institute for Molecular Science (LIMS), La Trobe University, Australia
2. La Trobe University Bioimaging platform, La Trobe University, Australia
3. Centre for Cardiovascular Biology & Disease Research, La Trobe Institute for Molecular Science (LIMS), La Trobe University
1. **Supplementary Methods**

**Solvent-doped alginate cell viability assay methodology**

For solvent doped cell viability assays the 5 mg/ml alginate hydrogels were supplemented with 0.05% (v/v) DMSO or HFIP to test if the solvent present in the hybrid materials was toxic to cells. The solvent doped hydrogels were bioprinted into 96 well plates and crosslinked using 50 mM CaCl_2_ (Chem-Supply). Scaffolds were washed six times with DMEM (Gibco), supplemented with 1% penicillin and streptomycin (Gibco), 10% foetal bovine serum (FBS) (Gibco) and 2 mM Glutamine (Gibco) before leaving in the media overnight. Murine RAW264.7 cells were then seeded on the scaffolds at 250,000 cells/mL. Resazurin-based AlamarBlue-HS (Sigma-Aldrich) dye was added 2 hours before the plate read and incubated at 37°C 10% CO_2_. Post incubation fluorescence spectra was recorded on days 0, 1, 3 and 5 using a CLARIOstar plate reader (BMG Labtech) (excitation 560 nm, emission 590 nm) at 25°C using the top optic. Scaffold-only controls were used to subtract background fluorescence readings from samples. Statistical analysis was performed in GraphPad Prism using two-way ANOVA (with Tukey comparison).

**Mass Spectrometry**

Fmoc-FF peptides were solubilised in DMSO and HFIP at 100 mg/mL then diluted to 10-100 μM using LCMS grade 100% isopropanol in glass vials. The same isopropanol was used as a blank before running the Fmoc-FF solutions. The samples were analysed at 10 uM and 100 μM concentration by direct infusion electrospray ionisation-mass spectrometry (ESI-MS) using an Agilent 6530 QTOF mass-spectrometer.  The mass-analyser was set to 50-1700m/z range in positive ionisation mode, with multiple fragmentation voltages (20 V,50 V,100 V).  Spectra were processed and exported using Agilent Masshunter Qualitative browser software.

2.
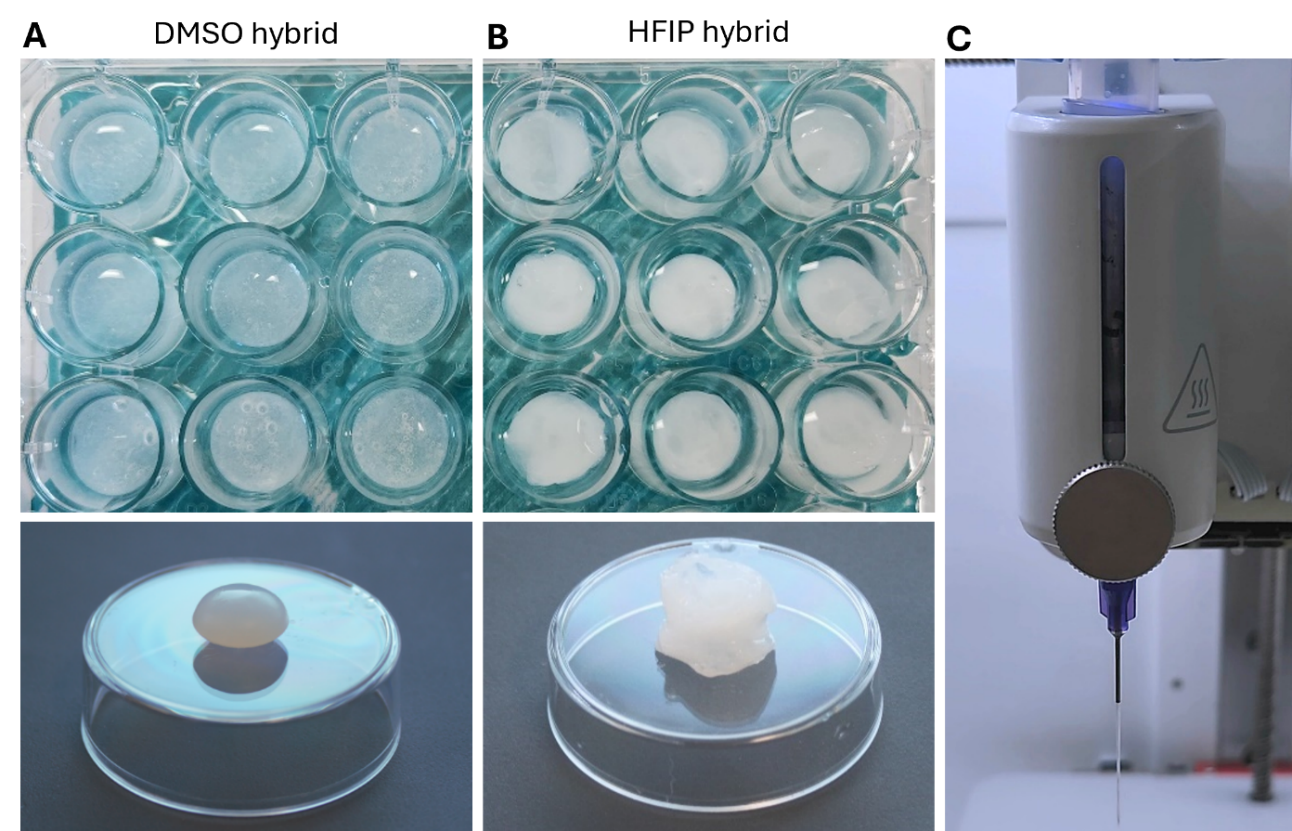
**Supplementary Figures**

**Supplementary figure 1:** Image of crosslinked DMSO hybrid (a) and HFIP hybrid (b) scaffold indicating uneven shape of HFIP hybrids and demonstration of continuous filament of hydrogel during printing (c).

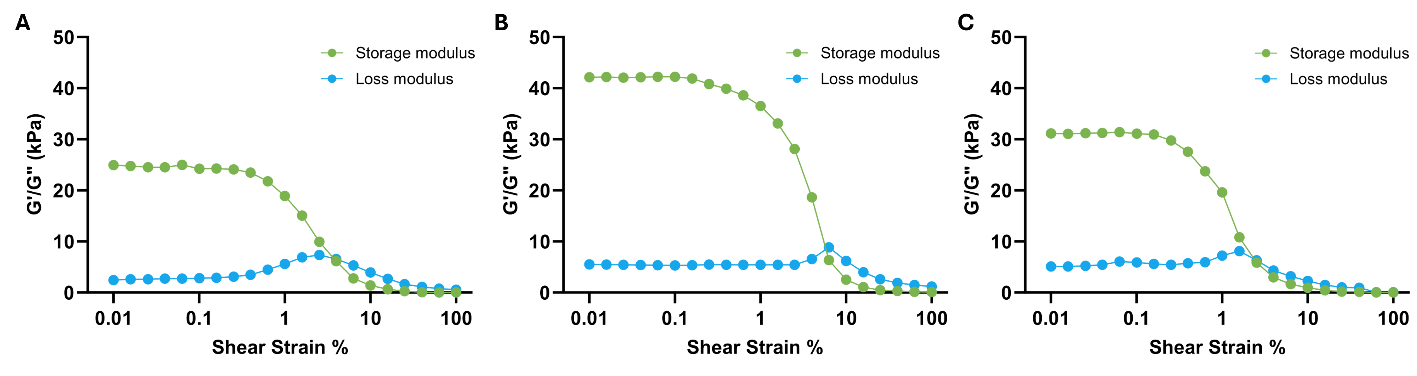

**Supplementary figure 2:** Amplitude sweeps of alginate (a), hybrid DMSO (b) and hybrid HFIP (c) scaffolds from 0.01-100% shear strain. Note the linear viscoelastic region (LVR) from 0.01-0.1% shear strain.

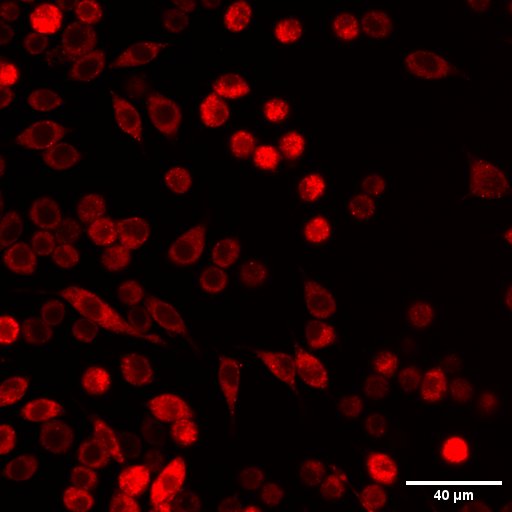

**Supplementary figure 3:** Laser scanning confocal microscopy image of RAW264.7 macrophages stained with CellTracker Orange 24 hours post-seeding on microscopy chamber slide. Scale bar shows 40 µm.

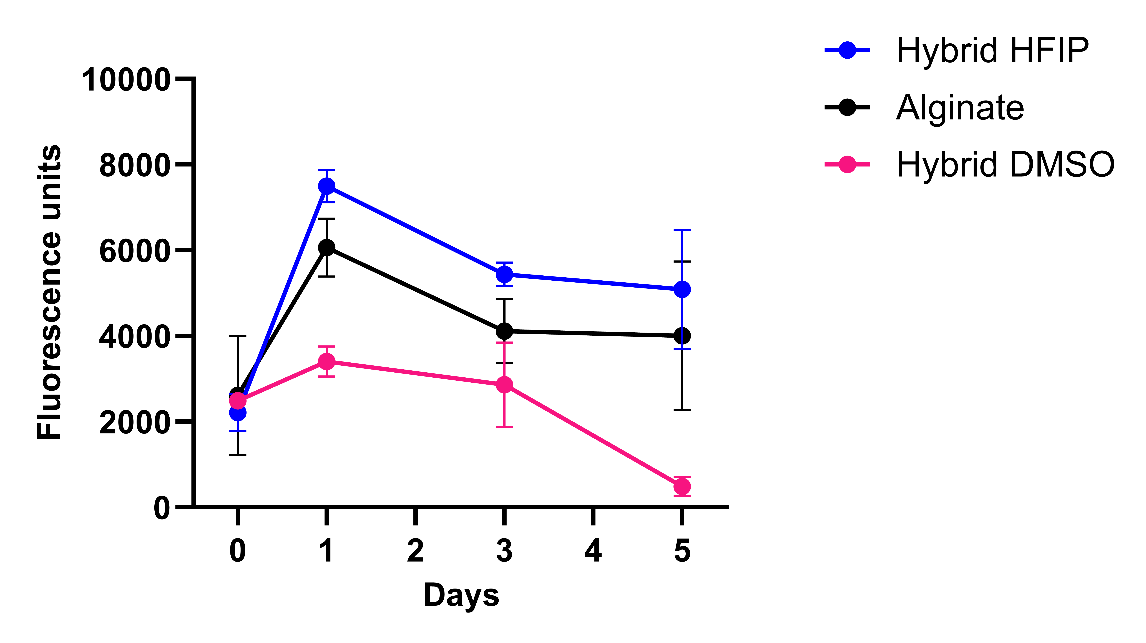
**Supplementary figure 4:** Viability of RAW264.7 macrophages seeded on top of alginate and hybrid hydrogels over 5 days. Data points represent the mean of 3 independent biological replicates (n=3) and error bars show the standard deviation of the mean.

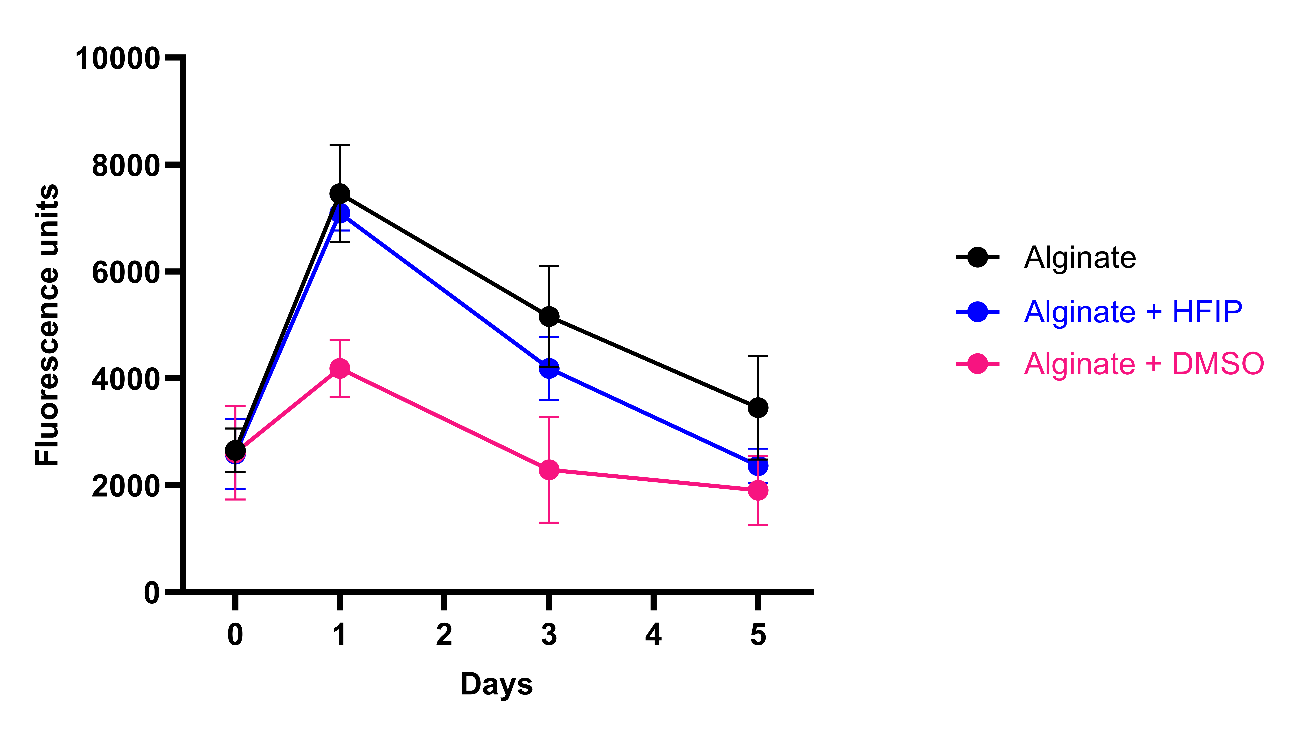

**Supplementary figure 5:** Viability of RAW264.7 macrophages seeded on top of 5% alginate scaffolds supplemented with 0.05 % DMSO and HFIP. Data points represent the mean of 3 independent biological replicates (n=3) and error bars show the standard deviation of the mean.

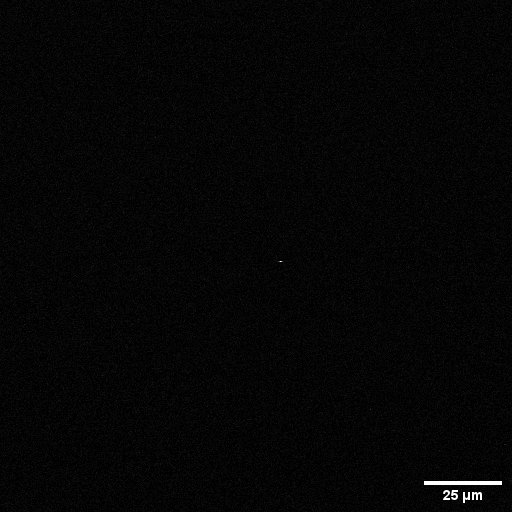

**Supplementary figure 6**: Laser scanning confocal microscopy image of alginate only scaffolds stained with Thioflavin T dye. No fluorescence detected demonstrating alginate does not bind Thioflavin T dye. Scale bar represents 25 µm.

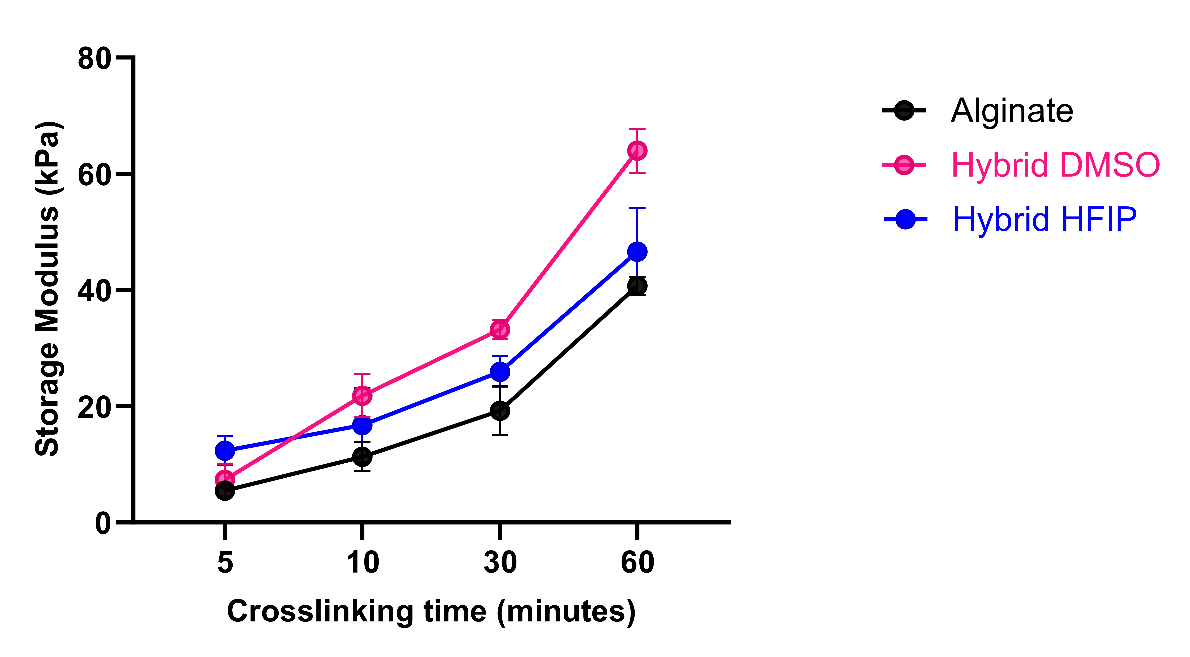

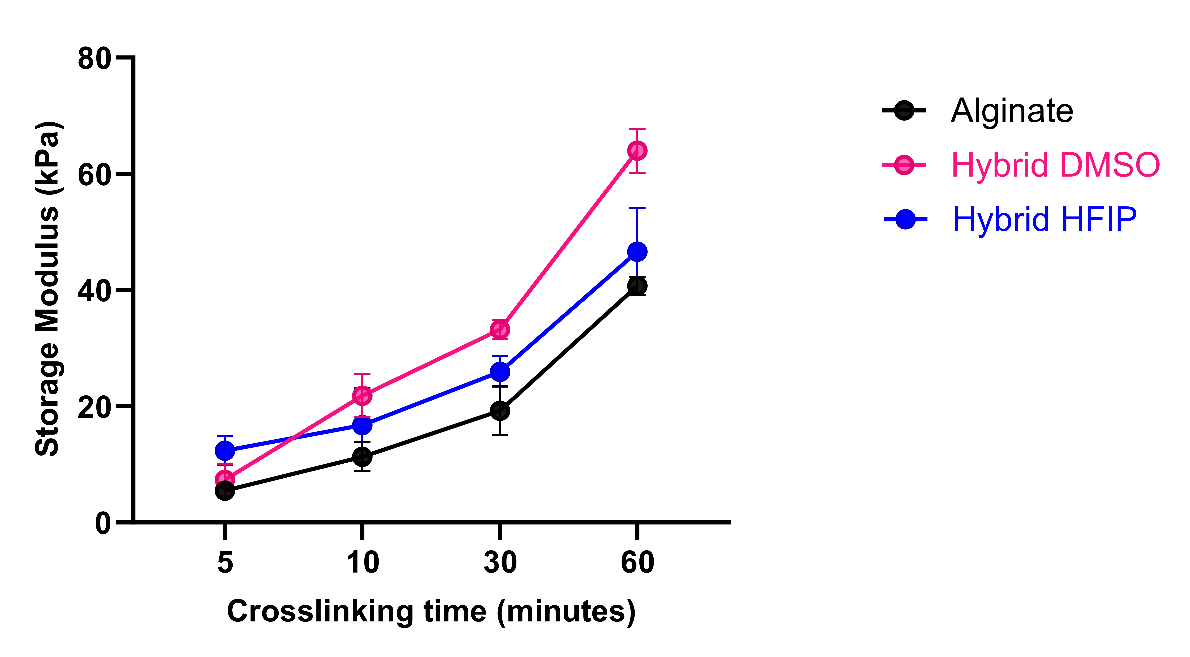

**Supplementary figure 7**: The effect of crosslinking time on the storage modulus of alginate and hybrid scaffolds at 0.1 % shear strain. Data points represent the mean of 3 independent biological replicates (n=3) and error bars show the standard deviation of the mean.

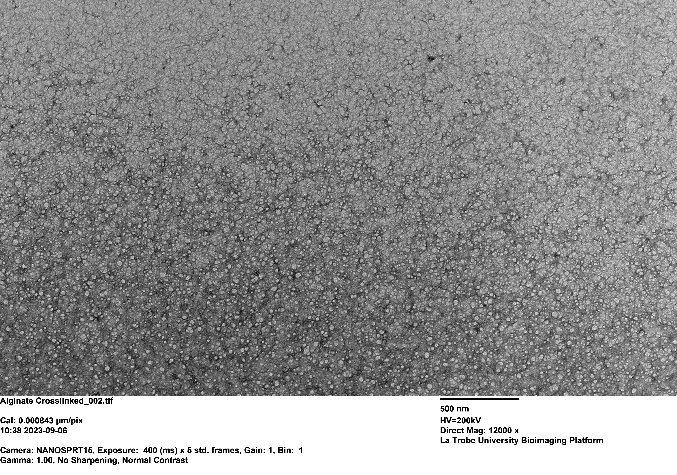

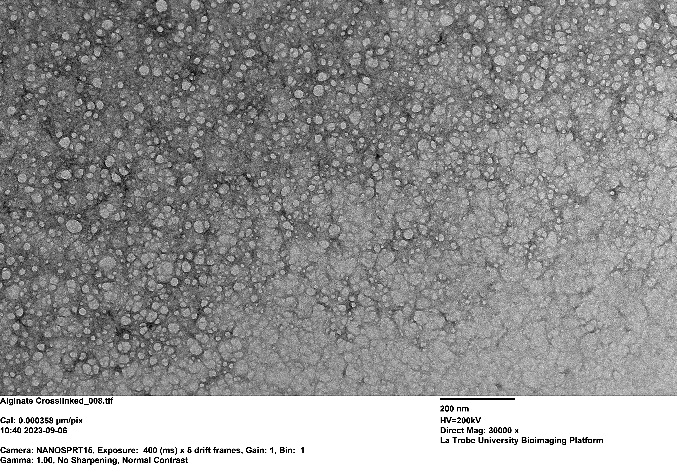

**Supplementary figure 8:** Transmission electron microscopy images of alginate-only samples

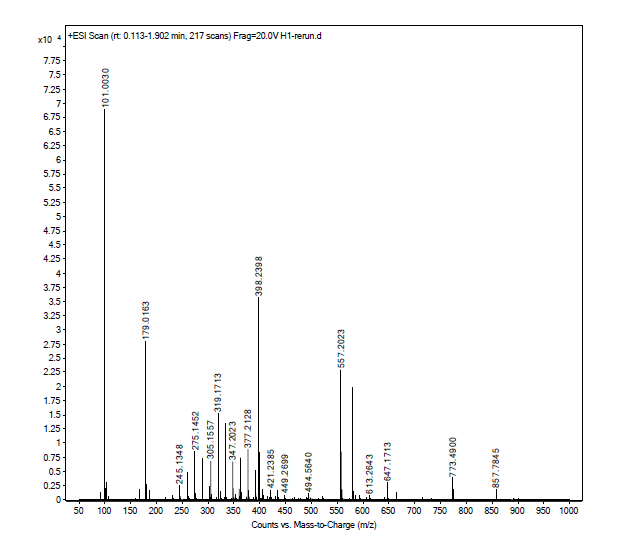

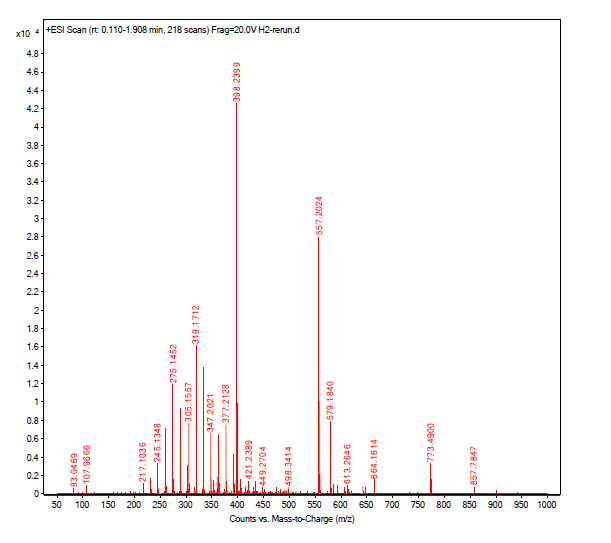

**Supplementary figure 9:** Mass spectrosocpy of Fmoc-FF peptides solubilised in DMSO (left) and HFIP (right) diluted to 100 μM with isopropanol. Results show no cleavage of Phe-Phe or Fmoc groups
